# The orientation of a membrane probe from structural analysis by enhanced Raman scattering

**DOI:** 10.1101/572529

**Authors:** Hannah J. Hughes, Steven M. E. Demers, Aobo Zhang, Jason H. Hafner

## Abstract

Small fluorescent molecules are widely used as probes of biomembranes. Different probes optically indicate membrane properties such as the lipid phase, thickness, viscosity, and electrical potential. The detailed molecular mechanisms behind probe signals are not well understood, in part due to the lack of tools to determine probe position and orientation in the membrane. Optical measurements on aligned biomembranes and lipid bilayers provide some degree of orientational information based on anisotropy in absorption, fluorescence, or nonlinear optical properties. These methods typically find the polar tilt angle between the membrane normal and the long axis of the molecule. Here we show that solution-phase surface enhanced Raman scattering (SERS) spectra of lipid membranes on gold nanorods can be used to determine molecular orientation of molecules within the membrane. The voltage sensitive dye 4-(2-(6-(dibutylamino)-2-naphthalenyl)ethenyl)-1-(3-sulfopropyl)-hydroxide, known as di-4-ANEPPS, is studied. Through the analysis of several peaks in the SERS spectrum, the polar angle from the membrane normal is found to be 63°, and the roll angle around the long axis of the molecule to be 305° from the original orientation. This structural analysis method could help elucidate the meaning of fluorescent membrane probe signals, and how they are affected by different lipid compositions.

## INTRODUCTION

Small molecule fluorescent probes are important tools for understanding structure and properties of biomembranes. These probes may signal specific membrane properties such as viscosity, permeability, lipid phase, or electric potential.^1-6^ Hundreds of membrane probes are available with molecular structures optimized for their intended purpose. For some probe methods, fluorophores are incorporated into the molecular structures of lipids or sterols. This yields fluorescent probes that can, for example, selectively partition into liquidordered (L_o_) or liquid-disordered (L_d_) phases of membranes, providing contrast to image lipid rafts and domains.^2^ Another approach is for fluorescent molecules to partition into the lipid membrane to generate signals. These probes may also serve as contrast agents based on how they partition, but their specific molecular properties can also be exploited. For example, fluorescent molecular rotors have rotational degrees freedom that affect their fluorescent properties and are also influenced by membrane viscosity.^7, 8^ These probes allow the detection and imaging of local rheological properties in a membrane. Sometimes the probe is designed to have a lipophilic exterior to enter the membrane, yet is also charged to drive passage through the membrane due to a transmembrane potential.^9^ However, the biophysical and photophysical properties of most probes are not entirely understood. The probe’s position and orientation in the membrane, the exact meaning of a probe’s signal, how that signal might be affected by membrane phase and composition, and how the probe may alter the inherent membrane properties are important open questions. ^10-14^

Here we focus on a class of “voltage sensitive dyes” which report on the electric fields inside membranes through changes in their fluorescent properties.^15, 16^ These dyes are based on aminostyryl pyridinium chromophores with alkane chains at one end and hydrophilic groups at the other end to orient the probe within the membrane. The strong electric field of the membrane changes the probe’s excitation and emission peaks. By carefully choosing excitation and detection wavelengths, the spectral shifts can be measured ratiometrically, independent of intensity fluctuations. While several effects could perturb fluorescence properties, such as probe reorientation or binding to other membrane components, the speed of response suggests that these probes work by a Stark shift mechanism. Recent work on TDDFT and MD supports this view, finding no effect due to intermolecular interactions.^17^

Lipid membranes create a complex electrostatic environment usually characterized by three potential differences: the surface potential (between the membrane surface and bulk solvent), the transmembrane potential (across the membrane, from one surface to the other) and the dipole potential (an internal potential barrier due to the arrangement of molecular dipoles).^18^ Since membrane probes are designed to sit inside the membrane, they are not generally affected by surface potential. The transmembrane potential controls signal transduction and other critical biomembrane processes. It creates an electric field inside the membrane and that can be studied in living cells with sub-second resolution with voltage sensitive membrane probes. The dipole potential is also internal to the membrane. It creates a much larger electric field that is highly localized to the interface between the hydrophobic and hydrophilic layers.^19^ The dipole potential is thought to affect internal membrane structure and therefore possibly influence signaling, but this is largely speculation due to the limited structural information available. Some probes, like di-4-ANEPPS studied here, have been shown to strongly respond to the dipole potential in addition to the transmembrane potential.^16^

The voltage sensitive probe’s response to the dipole potential depends on many factors: the alignment between the probe dipole moment and the membrane electric field, how the dipole direction changes upon excitation, the dipole direction relative to the probe structure (usually assumed to be the long axis), the orientation of the probe in the membrane, and the depth of the probe in the membrane.^20^ None of those parameters are easily determined given the difficulty of solving structures in lipid membranes.^21-25^ The molecular orientation of di-8-ANEPPS and di-4-ANEPPS have been studied with polarized fluorescence and second harmonic generation (SHG), as well as linear dichroism on aligned lipid membranes.^13, 26-29^ These measurements give a range of conflicting answer for the probe tilt from the membrane normal. Here we apply a method we recently introduced called structural analysis by enhanced Raman scattering (SABERS) to study the probe orientation.^30^ It combines surface enhanced Raman scattering (SERS), normal Raman scattering, electromagnetic fields calculated by the finite element method (FEM) and Raman tensors from time dependent density functional theory (TDDFT).

## MATERIALS AND METHODS

### Sample Preparation

The lipids 1,2-dioleoyl-sn-glycero-3-phosphocholine (DOPC) and 1,2-dioleoyl-sn-glycero-3-phospho-(1’-rac-glycerol) sodium salt (DOPG) were purchased from Avanti and combined at a 9:1 DOPC:DOPG molar ratio in chloroform. The lipids were dried down under a stream of nitrogen and placed under vacuum for 30 minutes. The probe pyridinium, 4-(2-(6-(dibutylamino)-2-naphthalenyl)ethenyl)-1-(3-sulfopropyl)-,hydroxide (di-4-ANEPPS) was purchased from Invitrogen. It was dissolved in ethanol and added to the dried lipid film at a 12:1 lipid:probe molar ratio. The solution was dried under a stream of nitrogen, and then hydrated with DI water to form multilamellar vesicles at a concentration of 10 mg/mL. To form small unilamellar vesicles (SUVs) the solution was water bath sonicated until the solution went from turbid to clear. Raman scattering experiments were recorded from these 10 mg/mL SUV solutions. For SERS experiments, gold nanorods were purchased from Nanopartz (nominally 50 nm diameter, 150 nm length, 800 nm peak absorbance, Lot #12H211). As received the nanorods were suspended in cetyltrimethylammonium bromide (CTAB) surfactant. Nanorod suspensions were pelleted by centrifugation at 1360 RCF for 15 min, and the supernatant was discarded. The nanorod pellet was then resuspended in 200 μL of phospholipid SUV prepared as described above at a concentration of 10 mg/mL. The nanorod/lipid solution was then bath sonicated for 5 min. The centrifugation and sonication were then repeated several times until the CTAB headgroup was not detectable by SERS analysis.^31^

### Gold Nanorod Characterization

3 μL of gold nanorod solution was dried onto a carbon coated grid and rinsed with water to minimize surfactant coating. The grids were imaged with a JEOL 1230 high contrast transmission electron microscope (TEM) at 80 keV.

### Raman and SERS Measurements

All spectra were recorded with a custom Raman microspectrometer. The excitation source is a stabilized diode laser (Ondax) at 785 nm wavelength and 80 mW power. The beam is further filtered by a volume holographic grating, and the power is adjusted with a linear variable neutral density filter. The beam is brought into the imaging system via a dichroic mirror. The beam is focused into the sample by a near-infrared corrected 40X/0.5 NA microscope objective (IR LCPlan N, Olympus). The samples are held in 2×2 mm capillaries with 0.1 mm thick walls (Vitrotube). The objective focusses the beam spot into the sample 50 μm past the glass/solution interface. Scattered light is collected by the objective, then passes through the dichroic mirror and through a notch filter. It is then focused with a second near-infrared corrected objective (5x/0.1 LMPlan N, Olympus) onto the entrance slit of a spectrograph (IsoPlane SCT 320, Princeton Instruments). The spectrum is recorded on a front-illuminated open electrode CCD camera (Pixis 256E, Princeton Instruments).

### Computational Methods

The electromagnetic near field surrounding the gold nanorods was calculated by solving Maxwell’s equations with the Finite Element Method (FEM) in COMSOL Multiphysics. The nanorod was constructed as a cylinder with hemispherical endcaps and given the dielectric constant of gold according to published values.^32^ The surrounding dielectric was that of water. The nanorod structure was also surrounded by a perfectly matched layer (PML). To simulate the relevant near field, an electromagnetic plane wave was incident on the nanorod and polarized along its length.

The di-4-ANEPPS derived polarizability tensor was calculated by TDDFT using the Amsterdam Density Functional package from Software for Chemistry & Materials (ADF2016).^33-35^ The geometry was optimized using the Becke-Perdew exchange-correlation potential under the generalized gradient approximation (GGA-BP) with a triple zeta basis set with two polarization functions (TZ2P), a large frozen core, and scalar ZORA relativistic correction.^36-39^ The numeric quality was set to “good”. Once the geometry was optimized, a vibrational Raman optical activity (VROA) calculation was carried out for 785 nm excitation with numerical frequencies and two-point numerical differentiation to find the derived polarizability tensor α for each mode.^40-45^

## RESULTS AND DISCUSSION

SABERS determines the orientation and position of a molecule near a nanoparticle based on the relative strength of SERS peaks caused by the electromagnetic near-field of the nanoparticle.^30^ Gold nanorods were used for this study because they are monodisperse, tunable to the excitation wavelength, and stable in solution. The gold nanorods used here were analyzed by TEM (Fig. 1a) which found an average rod diameter of 58 nm and a length of 154 nm. That geometry was used to calculate the far-field extinction and the electromagnetic near-field at frequencies ranging from the visible to near IR. The calculations were in good agreement with measured extinction spectra (Fig. 1b). The calculated electromagnetic near-field (Fig. 1c) decays within ca. 30 nm and is normal to the gold nanorod surface. The field decay is calculated as a function of distance from the gold surface for all Raman shifts (Fig. 1d). In the SERS analysis described below, we assume a normal electric field with a field strength and decay equivalent to the calculation result.

**Table.**
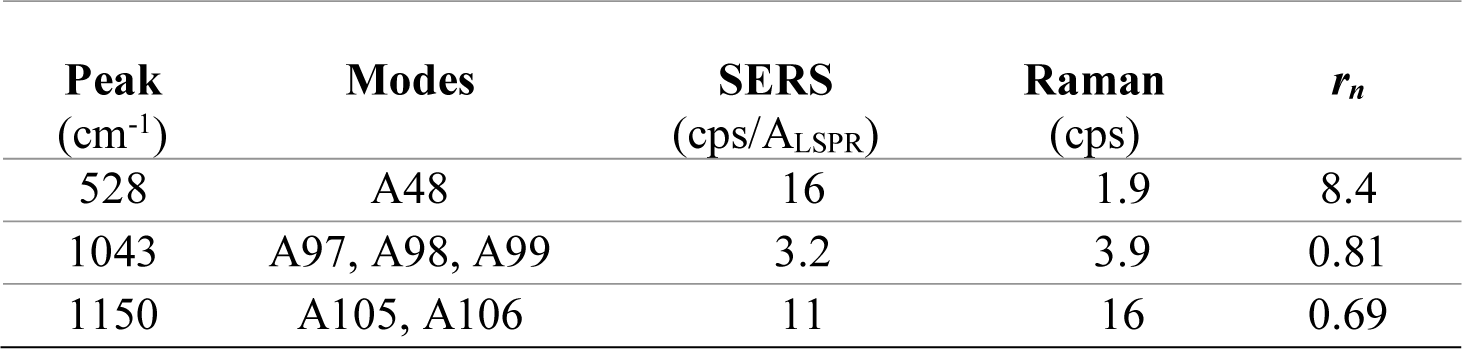
Spectral data for the three Raman and SERS peaks used in this study. The peak wavenumbers are the experimentally observed values. The mode labels correspond to a list of all modes with increasing energy according to TDDFT.

| Peak<br>(cm<sup>-1</sup>) | Modes | SERS<br>(cps/A<sub>LSPR</sub>) | Raman<br>(cps) | $r_n$ |
| --- | --- | --- | --- | --- |
| 528 | A48 | 16 | 1.9 | 8.4 |
| 1043 | A97, A98, A99 | 3.2 | 3.9 | 0.81 |
| 1150 | A105, A106 | 11 | 16 | 0.69 |

**Fig. 1.**
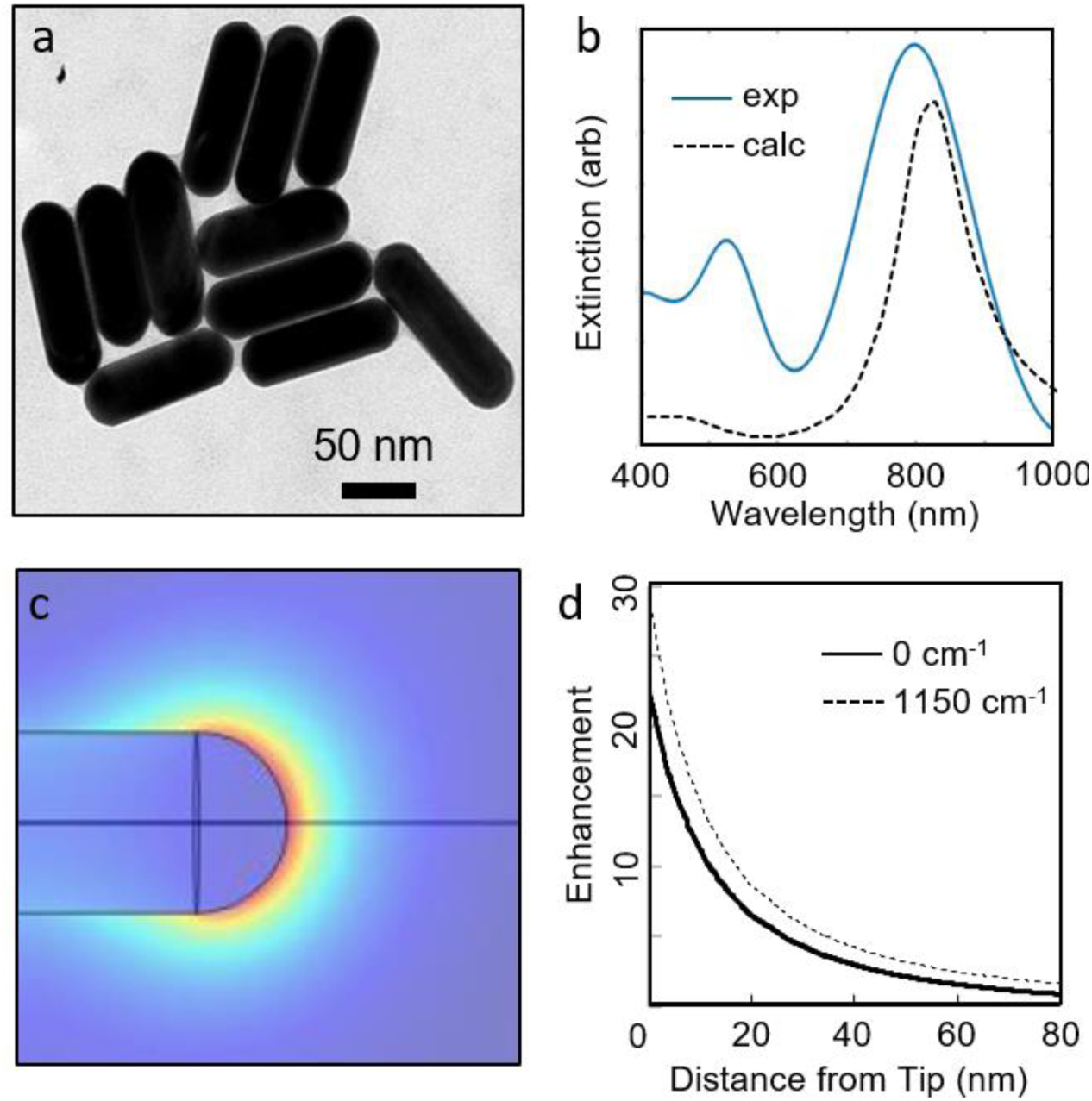
Gold nanorod characterization. The structure was determined from a) TEM image of the gold nanorods, and the average size and shape were used to b) calculate the (dashed) extinction spectrum and compare it to the measured absorption (solid). The calculation also generates c) a map of the electromagnetic field enhancement and d) the field decay from the nanorod tip at the excitation wavelength (785 nm, solid) and 1150 cm^−1^ shifted wavelength (862 nm, dashed) that are used in the analysis.

SABERS relies on ratiometric analysis of SERS and Raman peaks recorded under identical conditions. The wavelength shifted scattering spectra of lipid SUV’s with di-4-ANEPPS without nanorods (unenhanced) and with nanorods (enhanced) are compared (Fig. 2). Given the large electromagnetic enhancement at the end of the gold nanorods (∼10^5^), one might expect the two spectra to simply represent the Raman and SERS signals. However, in a nanorod/lipid suspension, the concentration of lipid vesicles not associated with nanorods is many orders of magnitude higher than the concentration of lipids on gold nanorods. In our experiments that concentration difference and the surface enhancement offset so that we get similar strength SERS and Raman contributions when nanorods are in the solution. A simple way forward is to subtract the unenhanced signal from the enhanced signal to isolate the SERS component, but we find that nanorod absorption can make this unreliable. We therefore record spectra at several nanorod concentrations. Plots of peak intensity versus nanorod concentration for three modes were corrected for absorption by the nanorod solution and fit by linear regression (Fig. 3). This measurement isolates the SERS component as the slope of the line, and the Raman component as the y-intercept. It also reduces error by providing these numbers from multiple experiments rather than just two (with and without nanorods).

**Fig. 2.**
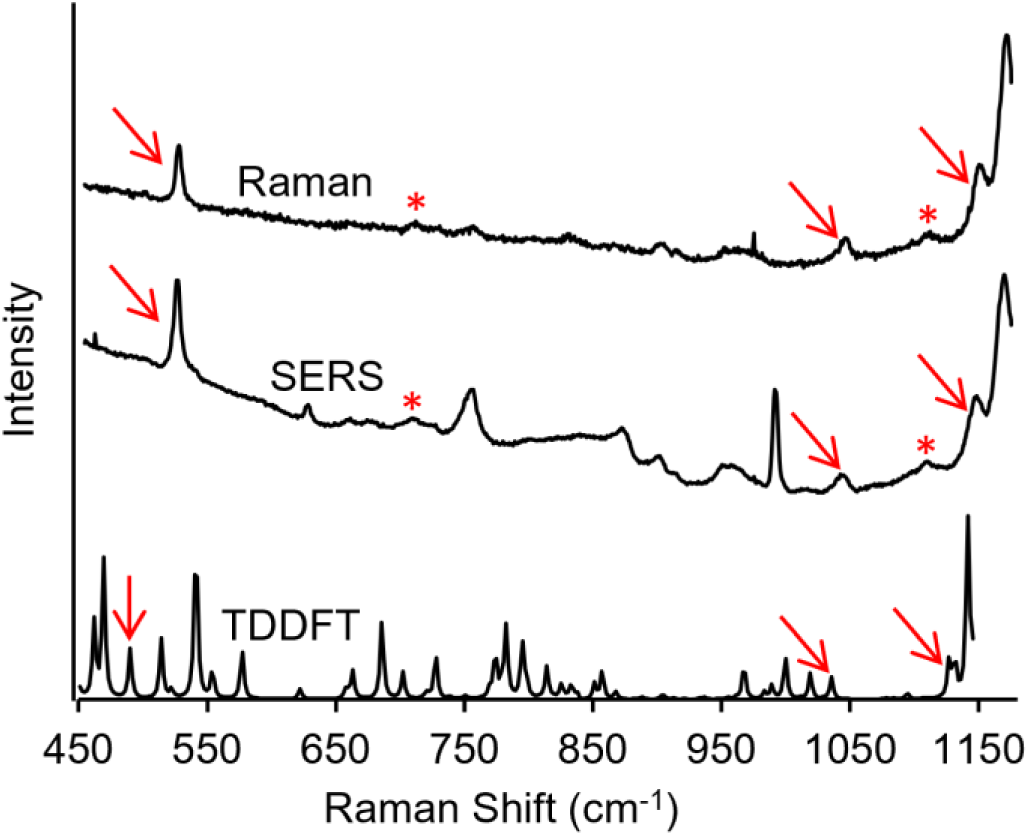
The experimental Raman and SERS spectra for di-4-ANEPPS in lipid vesicles, and the TDDFT calculated Raman spectrum. Asterisks indicate lipid modes at 718 cm^−1^ (symmetric choline stretch) and 1125 cm^−1^ (skeletal C-C). Arrows indicate peaks used in the current analysis and listed in the Table.

**Fig. 3.**
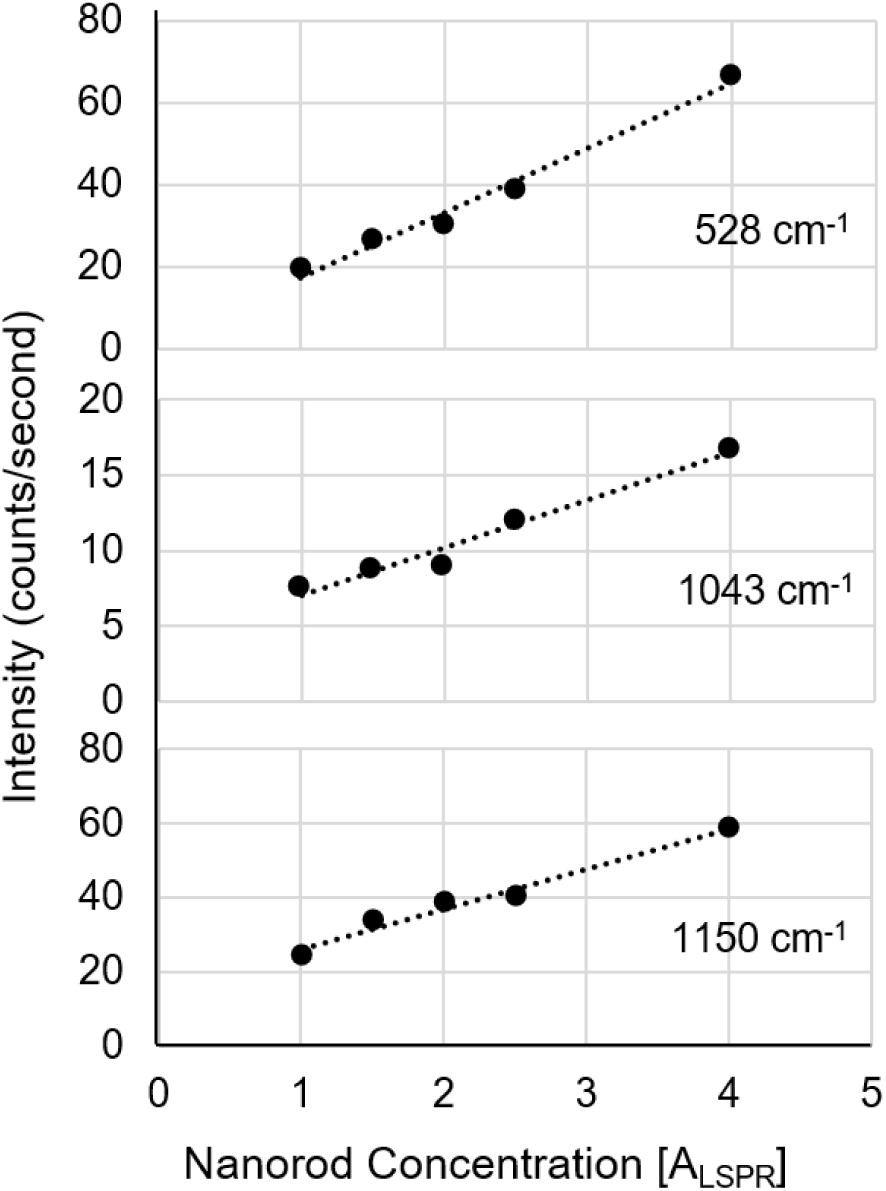
Spectral peak intensity as a function of gold nanorod concentration (in terms of plasmon resonance absorption maximum) for three peaks: a. 528 cm^−1^, b. 1043 cm^−1^, and c. 1150 cm^−1^. The slopes of these lines indicate the SERS signal, and the y-intercept indicates the Raman signal.

Several criteria are applied when choosing spectral peaks for analysis. First, the peak must be observed in spectra both with and without gold nanorods. Although “legitimate” modes could exist that are unobservable without enhancement, this criteria guards against spurious peaks related to irrelevant molecules adsorbed to the nanorods. Second, a plot of the peak intensity versus nanorod concentration must be linear to confirm that it is associated with the gold nanorods, as in Fig. 3. Third, the experimentally observed peak must also be identified among the modes calculated by TDDFT. Sometimes the match is clear, and sometimes trial and error is needed in the analysis (checking several nearby peaks to find which gives a reasonable result). Three peaks associated with di-4-ANEPPS passed these criteria and are highlighted with arrows in Fig. 2. The peak at 1150 cm^−1^ is easily identified in TDDFT based on its relative position to a stronger peak nearby. TDDFT predicts two modes (A105, A106) at that spectral location, so both were used in the analysis. A105 is primarily a stretch of the bond between the pyridinium nitrogen and the first carbon along the chain to the sulfate. A106 is similar. The peak at 1043 cm^−1^ is the last of a band of modes between 950 and 1050 cm^−1^ and corresponds to three skeletal modes of the three carbon chains (A97, A98, A99). Finally, the strong peak at 528 cm^−1^ corresponds to a mode involving deformations of the entire ring system (A48). Animations of the modes employed can be seen in the Supporting Information. Two peaks that correspond to lipid modes were also observed and are marked with an asterisk: the symmetric stretch of the choline headgroup at 718 cm^−1^ and the skeletal C-C vibrations at 1125 cm^−1^. However, they were not used for SABERS analysis due to their close proximity to other peaks (718 cm^−1^) and a lack of TDDFT analysis of the entire lipid molecule (1125 cm^−1^).

The analyzed peaks, assigned modes, and Raman and SERS intensities from Fig. 3 are summarized in the Table. SABERS bases the analysis of these data on the semiclassical description of Raman light scattering:

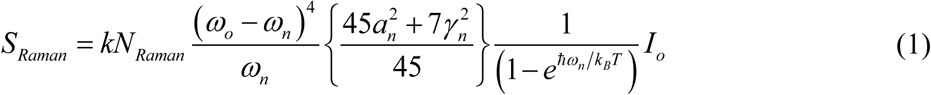

where *S*_*Raman*_ is the signal intensity in counts per seond.^46^ *I*_*o*_ is the excitation power of the laser, *ω*_*o*_ is the excitation frequency, *ω*_*n*_ is the Raman shifted frequency for the *n*^th^ normal mode, *k*_*B*_ is Boltzmann’s constant, *T* is the temperature, *α* and *γ* are Raman tensor invariants (average polarizability and average anisotropy, respectively), and *N*_*Raman*_ is the number of molecules in the beam spot. *k* absorbs all fundamental constants and instrumental parameters that convert the signal to counts/second. Three modifications are made for SERS. First, enhancement factors *E*_*n, o*_ and *E*_*n*,*R*_ are added to represent electromagnetic enhancement at the excitation and Raman shifted frequencies. Second, the orientation-averaged tensor invariants are replaced with *α* _*zz*_^2^, the Raman tensor component that corresponds to a normal electric field at the nanoparticle surface. Third, the number of molecules becomes *N*_*SERS*_, which represents the number of molecules in the enhanced region at the nanoparticle end:

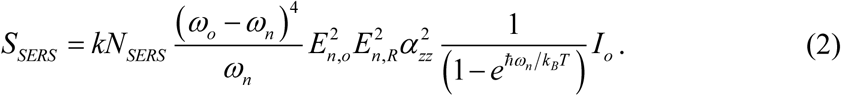

To remove the dependence on instrumental factors, such as the spectral efficiencies of the camera and spectrograph, the ratio, *r*_*n*_, between the SERS and Raman intensities is calculated, and included in the Table. Several factors cancel in the expression for *r*_*n*_:

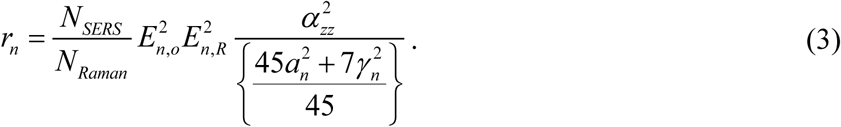

However, the ratio still includes the unknown number of molecules in the Raman and SERS measurement. The latter is especially difficult to determine and is usually the most significant unknown parameter in attempts to get quantitative information from SERS analysis.^47, 48^ We therefore take a second ratio between two peaks within the spectrum:

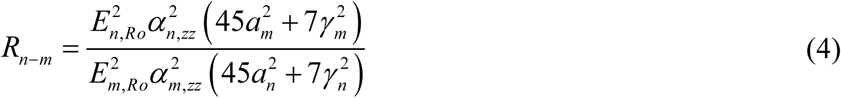

Note that this second ratio for two modes, *R*_*n-m*_, can be determined from the experimental values in the Table and calculated with field enhancements from FEM simulations and Raman tensor elements from TDDFT.

Theoretical *R*_*n-m*_ values are calculated for all rotations of the molecule relative to the nanorod surface (and therefore the direction of the electric field) by rotating the molecular structure and the Raman tensors. The rotations are defined in terms of a polar angle *θ* from the membrane normal, and the roll angle *φ* around the molecule’s long axis. Based on the initial orientation in Fig. 4a, *φ* (roll) rotations are first made around the *z*-axis and then *θ* (polar) rotations about the *y*-axis. Each vibration must also be assigned a specific distance from the nanorod surface for the field enhancement. We find a unique location for each mode from the individual vibration amplitudes of each atom in the molecule for that mode. The location is taken to be the mode’s amplitude-weighted averaged atomic position, which we refer to as the center of vibration:

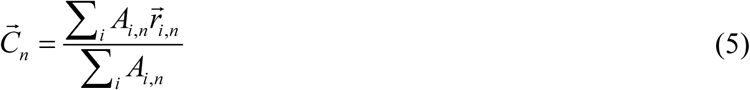

where ***C***_*n*_ is the center of vibration position vector for the *n*^th^ mode, *A*_*i*,*n*_ and ***r***_*i*,*n*_ are the vibration amplitudes and average positions, respectively, for the *i*^*th*^ atom in the *n*^*th*^ mode. The modes used in this analysis are illustrated in Fig. 4, where the eigenvectors for each mode are plotted at their center of vibration. For animations of each mode see the supporting information.

**Fig. 4.**
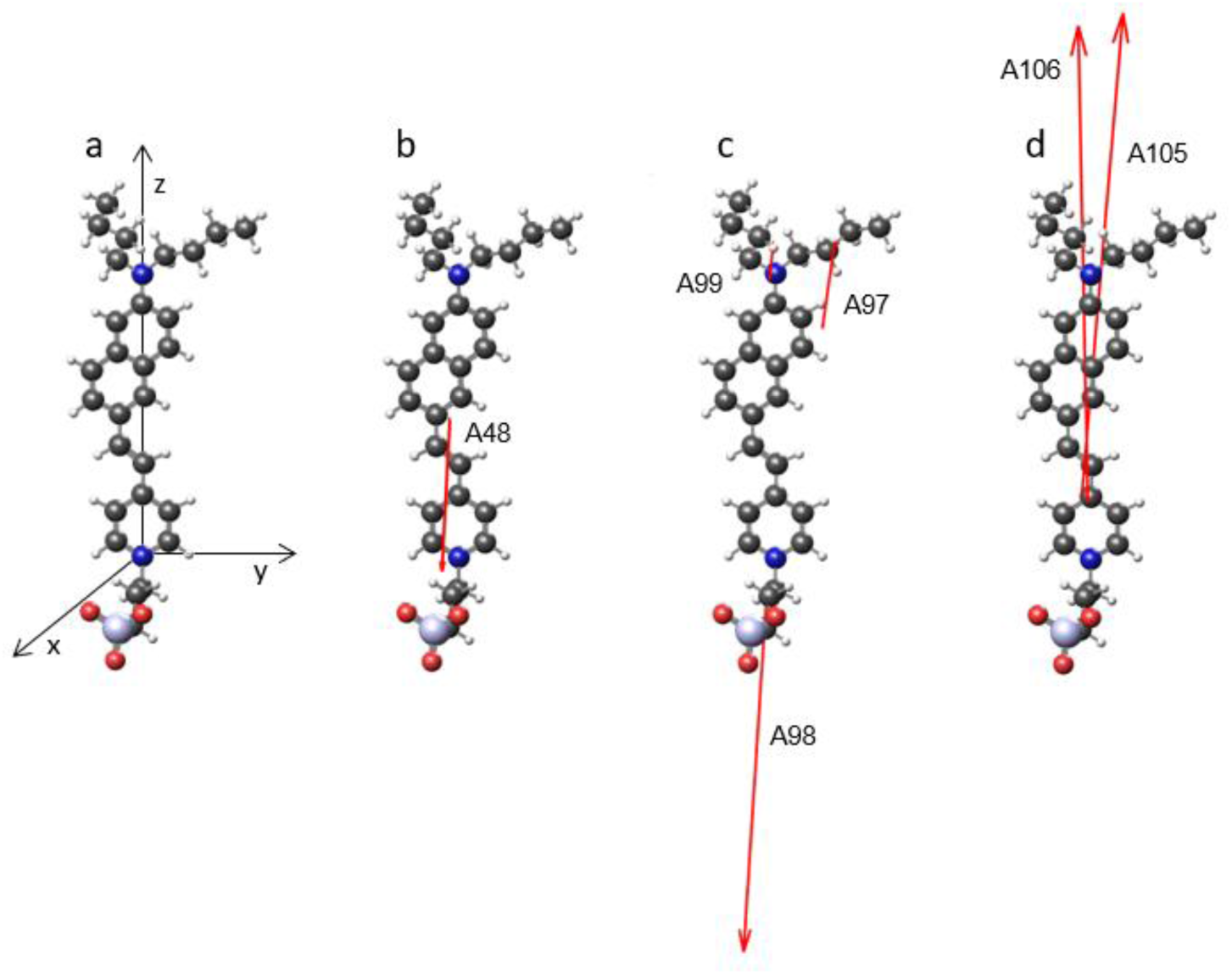
The molecular structure and Raman tensors. a) The original molecular orientation relative to Cartesian axes. The origin is on the pyridinium nitrogen atom and the z-axis passes through naphthylamine nitrogen atom. The pyridine and naphthylamine rings are in the y-z plane. The roll angle *ϕ* is adjusted by rotation around the z-axis, and the polar angle *θ* is adjusted by rotation around the y-axis. The eigenvectors for the modes that represents the peak at b) 528 cm^−1^, c) 1043 cm^−1^, and d) 1150 cm^−1^ are plotted to the same scale at their center of vibration.

The molecular orientation is determined by finding the best match between the experimentally measured value of *R*_*n-m*_ and the theoretically calculated values, using a similarity index defined as

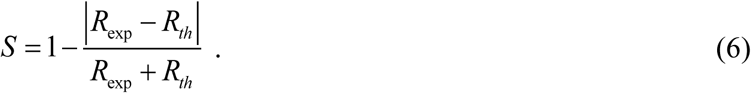

This index is 1 when the theoretical and experimental *R*’s match, and zero when they are very different. The similarity index is plotted at all theoretical molecular orientations to generate a map (Fig. 5) for each of the three pairs of peaks from the Table. The map for 528 cm^−1^ and 1043 cm^−1^ shows a band of high similarity values where the polar angle is above 45° and the roll angle is between 190° and 340°. The map for 528 cm^−1^ and 1150 cm^−1^ has high similarity for a somewhat lower polar angle and a wider range of roll angles. The similarity map for 1043 cm^−1^ and 1150 cm^−1^ has high values at almost all angles, indicating a lack of orientation dependence for that ratio. Each pair of modes has high similarity at a range of orientations, and therefore indicates a range of possible structures. However, the measurements were taken on a single sample that, presumably, had only one molecular orientation. We therefore multiply the maps to find the orientation for which *all* ratios have high similarity, which indicates the molecular orientation according to SABERS (Fig. 5d). According to this analysis, the di-4-ANEPPS orientation has a 66° polar angle and a 305° roll angle relative to the initial orientation (Fig. 4a).

**Fig. 5.**
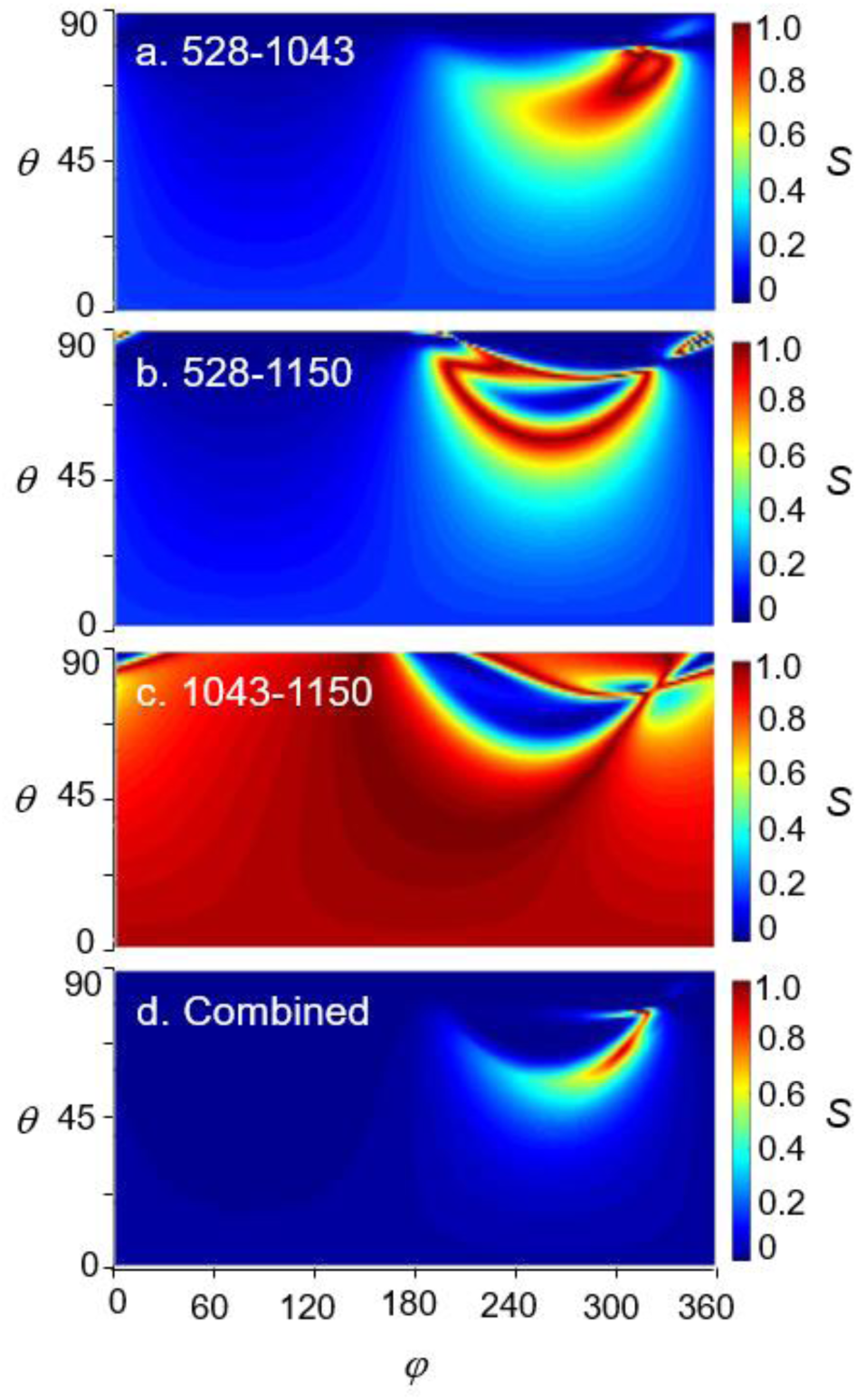
Similarity maps generated by calculating the similarity index at all polar and roll angles. Red regions indicate a similarity index close to one, while blue regions indicate a similarity index close to zero. a-c) The similarity maps for the indicated combination of modes calculated with eqns. 4 and 6. d) The combined similarity map obtained by multiplying the maps in a-c indicates a single orientation where the experimental and theoretical ratios are in agreement.

The molecular orientation of di-4-ANEPPS found here is depicted in Fig. 6, which shows the initial orientation, along with the roll and then polar rotations. During their initial development, di-4-ANEPPS and similar dyes were presumed to have their long axis normal to the membrane. Rather than a measurement, this was a putative structure based on the acyl chains and soluble groups at opposite ends of the molecule, and on the intended voltage sensitive function of the dye.^15, 49^ As these dyes have become widely used, optical methods have been applied to measure their orientation in membranes. Lambacher et al. analyzed the interferometric pattern of the fluorescence of di-8-ANEPPS in supported membranes on silicon substrates and found a 38° polar tilt angle.^26^ Ries et al. measured polarized second harmonic generation and two-photon fluorescence for di-8-ANEPPS in a black lipid membrane and found good agreement with a 36° tilt.^27^ However, Greeson et al. measured polarized fluorescence of di-8-ANEPPS from the membranes of living cells and found a tilt of 63° to the membrane normal, possibly pointing to the significant differences between pure lipid vesicles and natural biomembranes on probe orientation.^28^ Matson et al. measured the orientation of di-4-ANEPPS and di-8-ANEPPS from linear dichroism measurements on flow-aligned vesicles using retinoic acid as a reference and found smaller polar angles of 14° and 18° for each, respectively.^29^ Finally, Reeve et al. used one photon, two photon, and second harmonic generation imaging on GUV’s to measure both the polar tilt and its distribution. They found a polar angle of 52° for di-4-ANEPPS.^13^ These measurement results are rather widely varied, but likely for good reasons. One is the longer acyl chains of di-8 compared to di-4. Although the longer chains were designed to reduce the rate of flip-flop between leaflets, one would also expect the longer chain to affect the molecular orientation in the amphipathic membrane environment.^11, 12^ Another reason may be orientation sensitivity to different membrane components and conditions such as temperature, lipid bilayer phase, and thickness.^50^ Finally, there are significant assumptions and unknowns in these measurements, in including estimated tensor components and quantum efficiencies. Transition dipole moments were assumed to be along the molecule (between nitrogen on the amino and pyridinium groups), which is widely accepted. It was also assumed that emission dipole moments have the same direction as those for excitation, but experimental evidence suggests that is not always the case.^28^

**Fig. 6.**
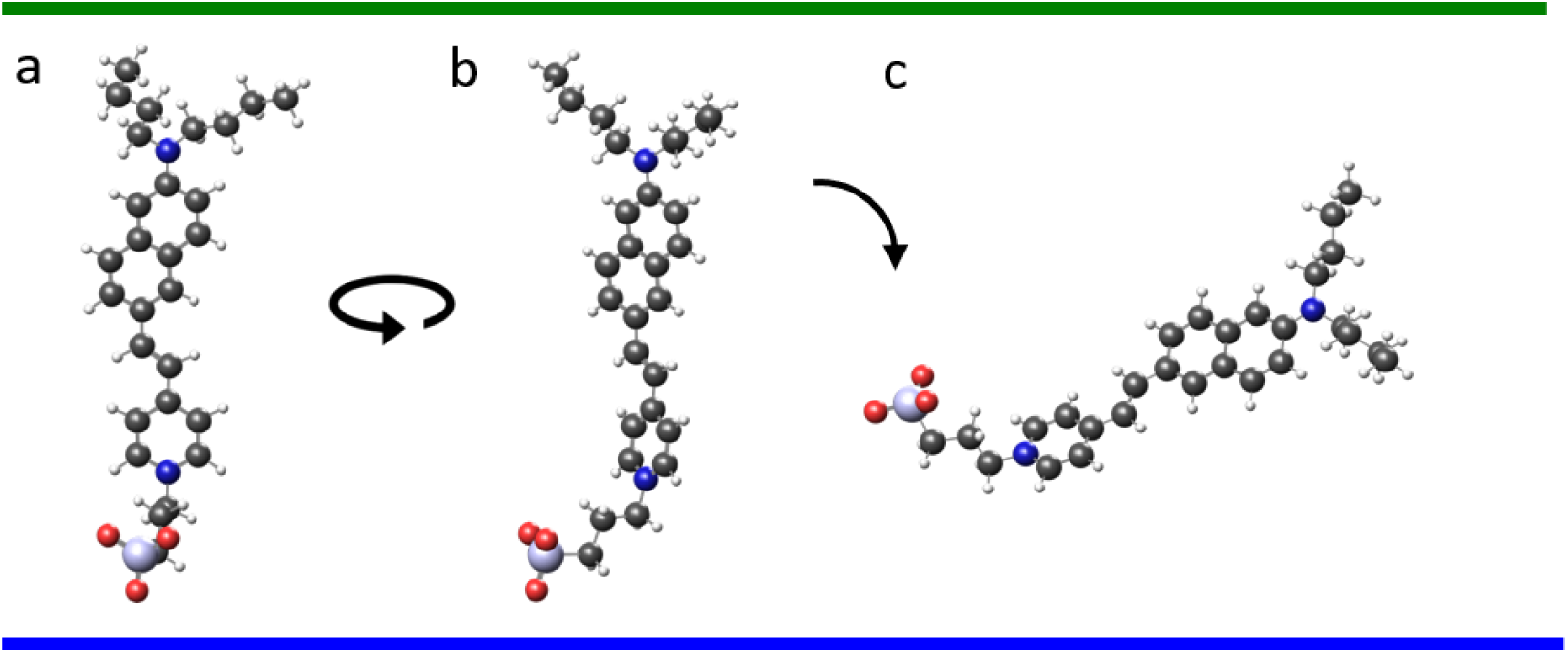
The molecular configuration of di-4-ANEPPS in the membrane. A. The initial orientation, B. the orientation after a *φ* = 305° rotation, and C. after a *θ* = 63° rotation. The horizontal lines represent the width of a single membrane leaflet.

The orientation reported here from SABERS for di-4-ANEPPS is at a large tilt angle (63°) and closest to the result by nonlinear microscopy (52°) on GUV’s.^13^ Note that the much smaller angle (14°) from linear dichroism is calibrated to measurements on retinol, which is presumed to be perfectly normal based on measurements on the retinol group in bacteriorhodopsin.^29^ It is possible that free retinol in a membrane is at a tilted angle (like all other optical probes studied to date), which would increase the estimated angles of di-4-ANEPPS and di-8-ANEPPS by that method.

A unique aspect of SABERS is that it finds the molecular roll angle in addition to the polar tilt within the bilayer since it is based on the alignment of the full polarizability tensors rather than just estimates of a property along the molecule. However, given that the bilayer is in the L_d_ state, it is not clear that the di-4-ANEPPS would take a specific roll orientation, or if it would spin around its long axis. Most of the measurements to date based on optical anisotropy cannot make this distinction, and one report pointed out that it affects interpretation of the structure.^27^ As presented above, SABERS indicates a static orientation for both polar and roll angles as the structure that matches the measured peak ratios. To model a rolling di-4-ANEPPS molecule, we ran the analysis by averaging the theoretical signal from two simultaneous orientations: the initial configuration (Fig. 4) and that orientation plus a 90° roll rotation. For more axially symmetric molecules like CTAB we have found this improves the analysis. However, the resulting similarity map show no signs of an orientation where the experimental and theoretical *R* values were in good agreement, thus indicating that our static solution (Fig. 6) is preferred. Future work will consider the effect of how small distributions of both polar and roll angles would affect the analysis.

In conclusion, a new structural analysis method based on SERS from lipid bilayers on gold nanorods, SABERS, has been applied to the voltage sensitive probe di-4-ANEPPS. Several di-4-ANEPPS spectral peaks were assigned to specific vibrational modes whose Raman tensor components were calculated by TDDFT. A double ratiometric analysis was employed to remove experimental unknowns so that both the tilt and roll angles of the di-4-ANEPPS in the membrane were determined. To gain further insight into these probes, TDDFT could also calculate the excited state geometry and the resulting change in dipole moment could be used to directly test the influence of electrochromism versus other possible phenomena like solvatochromism.^29^ Note that in our prior report on tryptophan, both the orientation and the z-position in the membrane were found by SABERS.^30^ Combined orientation and z-position information for voltage sensitive dyes could help clarify both the signaling mechanism and which membrane potentials they probe. Such information would also be beneficial to the new generation of dyes with increased sensitivity for in vivo imaging, such as photoactivatable voltage fluors and voltage sensitive dyes with tethered quenchers.^51, 52^

## Supporting information

A48 animation

A98 animation

A98 animation

A99 animation

A105 animation

A106 animation

## ACKNOWLEDGMENT

The authors acknowledge the Welch Foundation (grant C-1761) and National Science Foundation (award number 1709084) for supporting this research.

## REFERENCES

1. Kepczynski, M.; Rog, T., Functionalized lipids and surfactants for specific applications. Bba-Biomembranes 2016, 1858 (10), 2362–2379.

2. Klymchenko, A. S.; Kreder, R., Fluorescent Probes for Lipid Rafts: From Model Membranes to Living Cells. Chem Biol 2014, 21 (1), 97–113.

3. Sinkeldam, R. W.; Greco, N. J.; Tor, Y., Fluorescent Analogs of Biomolecular Building Blocks: Design, Properties, and Applications. Chem Rev 2010, 110 (5), 2579–2619.

4. Livanec, P. W.; Dunn, R. C., Single-Molecule Probes of Lipid Membrane Structure. Langmuir 2008, 24 (24), 14066–14073.

5. Munishkina, L. A.; Fink, A. L., Fluorescence as a method to reveal structures and membrane-interactions of amyloidogenic proteins. Bba-Biomembranes 2007, 1768 (8), 1862–1885.

6. Demchenko, A. P.; Mely, M.; Duportail, M.; Klymchenko, A. S., Monitoring Biophysical Properties of Lipid Membranes by Environment-Sensitive Fluorescent Probes. Biophys J 2009, 96 (9), 3461–3470.

7. Suhling, K., Twist and Probe-Fluorescent Molecular Rotors Image Escherichia coli Cell Membrane Viscosity. Biophys J 2016, 111 (7), 1337–1338.

8. Mika, J. T.; Thompson, A. J.; Dent, M. R.; Brooks, N. J.; Michiels, M.; Hofkens, M.; Kuimova, M. K., Measuring the Viscosity of the Escherichia coil Plasma Membrane Using Molecular Rotors. Biophys J 2016, 111 (7), 1528–1540.

9. Xu, M.; Zeng, Z. B.; Jiang, J. H.; Chang, Y. T.; Yuan, L., Discerning the Chemistry in Individual Organelles with Small-Molecule Fluorescent Probes. Angew Chem Int Edit 2016, 55 (44), 13658–13699.

10. Faller, R., Molecular modeling of lipid probes and their influence on the membrane. Bba-Biomembranes 2016, 1858 (10), 2353–2361.

11. Timr, M.; Bondar, M.; Cwiklik, M.; Stefl, M.; Hof, M.; Vazdar, M.; Lazar, M.; Jungwirth, P., Accurate Determination of the Orientational Distribution of a Fluorescent Molecule in a Phospholipid Membrane. J Phys Chem B 2014, 118 (4), 855–863.

12. Ferrand, M.; Gasecka, M.; Kress, M.; Wang, M.; Bioud, F. Z.; Duboisset, M.; Brasselet, S., Ultimate Use of Two-Photon Fluorescence Microscopy to Map Orientational Behavior of Fluorophores. Biophys J 2014, 106 (11), 2330–2339.

13. Reeve, J. E.; Corbett, A. D.; Boczarow, M.; Wilson, M.; Bayley, M.; Anderson, H. L., Probing the Orientational Distribution of Dyes in Membranes through Multiphoton Microscopy. Biophys J 2012, 103 (5), 907–917.

14. Skaug, M. J.; Longo, M. L.; Faller, R., The Impact of Texas Red on Lipid Bilayer Properties. J Phys Chem B 2011, 115 (26), 8500–8505.

15. Fluhler, M.; Burnham, V. G.; Loew, L. M., Spectra, Membrane-Binding, and Potentiometric Responses of New Charge Shift Probes. Biochemistry-Us 1985, 24 (21), 5749–5755.

16. Gross, M.; Bedlack, R. S.; Loew, L. M., Dual-Wavelength Ratiometric Fluorescence Measurement of the Membrane Dipole Potential. Biophys J 1994, 67 (1), 208–216.

17. Robinson, M.; Besley, N. A.; O’Shea, M.; Hirst, J. D., Di-8-ANEPPS Emission Spectra in Phospholipid/Cholesterol Membranes: A Theoretical Study. J Phys Chem B 2011, 115 (14), 4160–4167.

18. Clarke, R. J., The dipole potential of phospholipid membranes and methods for its detection. Adv Colloid Interfac 2001, 89, 263–281.

19. Yang, M.; Mayer, K. M.; Wickremasinghe, N. S.; Hafner, J. H., Probing the Lipid Membrane Dipole Potential by Atomic Force Microscopy. Biophys J 2008, 95 (11), 5193–5199.

20. Loew, L. M.; Scully, M.; Simpson, M.; Waggoner, A. S., Evidence for a Charge-Shift Electrochromic Mechanism in a Probe of Membrane-Potential. Nature 1979, 281 (5731), 497–499.

21. Sezgin, E., Super-resolution optical microscopy for studying membrane structure and dynamics. J. Phys.-Condes. Matter 2017, 29 (27).

22. Lee, T. H.; Hirst, D. J.; Kulkarni, M.; Del Borgo, M. P.; Aguilar, M. I., Exploring Molecular-Biomembrane Interactions with Surface Plasmon Resonance and Dual Polarization Interferometry Technology: Expanding the Spotlight onto Biomembrane Structure. Chem Rev 2018, 118 (11), 5392– 5487.

23. Rouck, J. E.; Krapf, J. E.; Roy, M.; Huff, H. C.; Das, A., Recent advances in nanodisc technology for membrane protein studies (2012-2017). Febs Lett 2017, 591 (14), 2057–2088.

24. Pluhackova, M.; Bockmann, R. A., Biomembranes in atomistic and coarse-grained simulations. J. Phys.-Condes. Matter 2015, 27 (32).

25. Malhotra, M.; Alder, N. N., Advances in the use of nanoscale bilayers to study membrane protein structure and function. Biotechnol Genet Eng 2014, 30 (1), 79–93.

26. Lambacher, M.; Fromherz, P., Orientation of hemicyanine dye in lipid membrane measured by fluorescence interferometry on a silicon chip. J Phys Chem B 2001, 105 (2), 343–346.

27. Ries, R. S.; Choi, M.; Blunck, M.; Bezanilla, M.; Heath, J. R., Black lipid membranes: visualizing the structure, dynamics, and substrate dependence of membranes. J Phys Chem B 2004, 108 (41), 16040–16049.

28. Greeson, J. N.; Raphael, R. M., Application of fluorescence polarization microscopy to measure fluorophore orientation in the outer hair cell plasma membrane. J Biomed Opt 2007, 12 (2).

29. Matson, M.; Carlsson, M.; Beke-Somfai, M.; Norden, B., Spectral Properties and Orientation of Voltage-Sensitive Dyes in Lipid Membranes. Langmuir 2012, 28 (29), 10808–10817.

30. Matthews, J. R.; Shirazinejad, C. R.; Isakson, G. A.; Demers, S. M. E.; Hafner, J. H., Structural Analysis by Enhanced Raman Scattering. Nano Lett 2017, 17 (4), 2172–2177.

31. Matthews, J. R.; Payne, C. M.; Hafner, J. H., Analysis of Phospholipid Bilayers on Gold Nanorods by Plasmon Resonance Sensing and Surface-Enhanced Raman Scattering. Langmuir 2015, 31 (36), 9893–9900.

32. Johnson, P. B.; Christy, R. W., Optical Constants of Noble Metals. Phys Rev B 1972, 6 (12), 4370–4379.

33. Baerends, E. J.; Ziegler, M.; Atkins, A. J.; Autschbach, M.; Bashford, M.; Baseggio, M.; Berces, M.; Bickelhaupt, F. M.; Bo, M.; Boerritger, P. M.; Cavallo, M.; Daul, M.; Chong, D. P.; Chulhai, D. V.; Deng, M.; Dickson, R. M.; Dieterich, J. M.; Ellis, D. E.; van Faassen, M.; Ghysels, M.; Giammona, M.; van Gisbergen, S. J. A.; Goez, M.; Gotz, A. W.; Gusarov, M.; Harris, F. E.; van den Hoek, M.; Hu, M.; Jacob, C. R.; Jacobsen, M.; Jensen, M.; Joubert, M.; Kaminski, J. W.; van Kessel, M.; Konig, M.; Kootstra, M.; Kovalenko, M.; Krykunov, M.; van Lenthe, M.; McCormack, D. A.; Michalak, M.; Mitoraj, M.; Morton, S. M.; Neugebauer, M.; Nicu, V. P.; Noodleman, M.; Osinga, V. P.; Patchkovskii, M.; Pavanello, M.; Peeples, C. A.; Philipsen, P. H. T.; Post, M.; Pye, C. C.; Ramanantoanina, M.; Ramos, M.; Ravenek, M.; Rodriguez, J. I.; Ros, M.; Ruger, M.; Schipper, P. R. T.; Schluns, M.; van Schoot, M.; Schreckenbach, M.; Seldenthuis, J. S.; Seth, M.; Snijders, J. G.; Sola, M.; Stener, M.; Swart, M.; Swerhone, M.; te Velde, M.; Tognetti, M.; Vernooijs, M.; Versluis, M.; Visscher, M.; Visser, M.; Wang, M.; Wesolowski, T. A.; van Wezenbeek, E. M.; Wiesenekker, M.; Wolff, S. K.; Woo, T. K.; Yakovlev, A. L., ADF2017, SCM, Theoretical Chemistry, Vrije Universiteit, Amsterdam, The Netherlands, https://www.scm.com.

34. Guerra, C. F.; Snijders, J. G.; te Velde, M.; Baerends, E. J., Towards an order-N DFT method. Theor Chem Acc 1998, 99 (6), 391–403.

35. te Velde, M.; Bickelhaupt, F. M.; Baerends, E. J.; Guerra, C. F.; Van Gisbergen, S. J. A.; Snijders, J. G.; Ziegler, T., Chemistry with ADF. J Comput Chem 2001, 22 (9), 931–967.

36. Vanlenthe, M.; Baerends, E. J.; Snijders, J. G., Relativistic Regular 2-Component Hamiltonians. J Chem Phys 1993, 99 (6), 4597–4610.

37. Vanlenthe, M.; Baerends, E. J.; Snijders, J. G., Relativistic Total-Energy Using Regular Approximations. J Chem Phys 1994, 101 (11), 9783–9792.

38. van Lenthe, M.; Ehlers, M.; Baerends, E. J., Geometry optimizations in the zero order regular approximation for relativistic effects. J Chem Phys 1999, 110 (18), 8943–8953.

39. Van Lenthe, M.; Baerends, E. J., Optimized slater-type basis sets for the elements 1-118. J Comput Chem 2003, 24 (9), 1142–1156.

40. Fan, L. Y.; Ziegler, T., Application of Density Functional Theory to Infrared-Absorption Intensity Calculations on Main Group Molecules. J Chem Phys 1992, 96 (12), 9005–9012.

41. Fan, L. Y.; Ziegler, T., Application of Density Functional Theory to Infrared-Absorption Intensity Calculations on Transition-Metal Carbonyls. J Phys Chem-Us 1992, 96 (17), 6937–6941.

42. Vangisbergen, S. J. A.; Snijders, J. G.; Baerends, E. J., A Density-Functional Theory Study of Frequency-Dependent Polarizabilities and Van-Der-Waals Dispersion Coefficients for Polyatomic-Molecules. J Chem Phys 1995, 103 (21), 9347–9354.

43. vanGisbergen, S. J. A.; Snijders, J. G.; Baerends, E. J., Application of time-dependent density functional response theory to Raman scattering. Chem Phys Lett 1996, 259 (5-6), 599-604.

44. van Gisbergen, S. J. A.; Snijders, J. G.; Baerends, E. J., Implementation of time-dependent density functional response equations. Comput Phys Commun 1999, 118 (2-3), 119–138.

45. Jensen, M.; Autschbach, M.; Krykunov, M.; Schatz, G. C., Resonance vibrational Raman optical activity: A time-dependent density functional theory approach. J Chem Phys 2007, 127 (13).

46. Long, D. A., Raman Spectroscopy. McGraw-Hill, Inc.: Great Britain, 1977; p 276.

47. Le Ru, E. C.; Blackie, M.; Meyer, M.; Etchegoin, P. G., Surface enhanced Raman scattering enhancement factors: a comprehensive study. J Phys Chem C 2007, 111 (37), 13794–13803.

48. Lee, M.; Anderson, L. J. E.; Payne, C. M.; Hafner, J. H., Structural Transition in the Surfactant Layer that Surrounds Gold Nanorods as Observed by Analytical Surface-Enhanced Raman Spectroscopy. Langmuir 2011, 27 (24), 14748–14756.

49. Loew, L. M., Design and Characterization of Electrochromic Membrane Probes. J Biochem Bioph Meth 1982, 6 (3), 243–260.

50. Wu, M.; Yeh, F. L.; Mao, M.; Chapman, E. R., Biophysical Characterization of Styryl Dye-Membrane Interactions. Biophys J 2009, 97 (1), 101–109.

51. Grenier, M.; Walker, A. S.; Miller, E. W., A Small-Molecule Photoactivatable Optical Sensor of Transmembrane Potential. J Am Chem Soc 2015, 137 (34), 10894–10897.

52. Yan, M.; Acker, C. D.; Loew, L. M., Tethered Bichromophoric Fluorophore Quencher Voltage Sensitive Dyes. Acs Sensors 2018, 3 (12), 2621–2628.

